# Laser capture microdissection-based single-cell RNA sequencing reveals optic nerve crush-related early mRNA alterations in retinal ganglion cells

**DOI:** 10.1101/2020.03.13.990077

**Authors:** Dongyan Pan, Mengqiao Xu, Xin Chang, Mao Xia, Yibin Fang, Yinghua Fu, Wei Shen, Yue Wang, Xiaodong Sun

## Abstract

Retinal ganglion cells (RGC) are the primary cell type injured in a variety of diseases of the optic nerve, and the early changes of RGC’s RNA profiling may be important to understand the mechanism of optic nerve injury and axon regeneration. Here we employed the optic nerve crush (ONC) model to explore early mRNA alterations in RGCs using laser capture microdissection (LCM) and single-cell RNA sequencing. We successfully established an optimal LCM protocol using 30 μm-thick retinal tissue sections mounted on glass slides and laser pressure catapulting (LPC) to collect RGCs and obtain high-quality RNA for single-cell sequencing. Based on our protocol, we identified 8744 differentially expressed genes that were involved in ONC-related early mRNA alterations in RGCs. Candidate genes included Atf3, Lgals3, LOC102551701, Plaur, Tmem140 and Maml1. The LCM-based single-cell RNA sequencing allowed new insights into the early mRNA changes in RGCs, highlighting new molecules associated with ONC.

## 1. Introduction

Retinal ganglion cell (RGC) death and axonal degeneration are common processes in optic nerve diseases, such as traumatic optic neuropathy and glaucoma. The optic nerve crush (ONC) model has proven useful for the study of RGC survival or death. RGC death commences within 5 to 6 days after ON injury, with more than 90% RGC loss after 14 days[1]. ONC-related mRNA changes have been documented in cell and animal models. However, RNA profiling studies focusing on early changes in RGCs after ONC are still rare.

Laser capture microdissection (LCM) allows the isolation of specific cell populations from complex tissues that can then be used for gene expression studies.[1] Single-cell RNA sequencing (scRNA-seq) can amplify a small amount of RNA (from at least one cell, approximately 10 pg) at the ng level to obtain the amount of nucleic acid needed for high-throughput sequencing. scRNA-seq can even reveal complex and rare cell populations, uncover regulatory relationships between genes, and track the trajectories of distinct cell lineages during development. Although studies involving 10x scRNA-seq of the retina have been reported recently,[2, 3] and their results have been valuable, there are still some shortcomings. This technique does not distinguish cells in advance; therefore, there is a large amount of sequencing data that cannot be referenced to target cells, which is not economical. In addition, no pathological sections are used as a reference, and it is unknown whether the samples are representative of the disease. Therefore, it may be of great significance to establish a more economical scRNA-seq method, especially for abnormal cells in pathological sections.

In the present study, we adopted an optimal protocol combined with LCM and scRNA-seq to examine the early changes in the RGC RNA profile in an ONC model. Our results also revealed optic nerve crush-related early mRNA alterations in retinal ganglion cells and identified several candidate genes for further studies.

## 2. Materials and methods

### 2.1. Animals and ONC model

Adult (150-200 g) male Wistar rats were included in the present study. The animals were housed in conditions with a constant room temperature (21–22 °C) and a 12 h light/12 h dark cycle and given free access to food and water. All experiments were performed according to the Guidance for the Care and Use of Laboratory Animals recommended by the National Institutes of Health and were approved by the Ethics Committee for Animal Experimentation of the Second Military Medical University.

To establish the ONC model, the rats were intraperitoneally anesthetized with 10% chloralhydrate (4 ml/kg), and intraorbital ONC was performed in the left eye as previously described.[4] Briefly, the optic nerve was surgically exposed under the superior orbital margin and crushed using an aneurysm clip (mini clip, temporary, Aesculap, Germany) 2 mm posterior to the lamina cribrosa for 8 s without damaging any small vessels. Sham surgery was performed in the right eye. Animals were sacrificed and perfused intracardially with PBS on day 3 postcrush.

### 2.2. Tissue Preparation

Eyes were removed corneas and lens and then embedded using optimal cutting temperature embedding medium (SAKURA Tissue-Tek® O.C.T. Compound) by rapid freezing under crushed dry ice and stored at −80°C.

Glass slides and PEN membrane-coated slides were disinfected with ultraviolet light, and the tissues were sectioned with a CM1950 cryostat microtome (Leica Microsystems Inc, Bannockburn, IL, http://www.leica-microsystems.com) at −22°C at a thickness of 20 µm or 30 μm.

### 2.3. Fixation and staining

The frozen sections were thawed at room temperature for 2 min and immersed immediately in 70% ethanol for 2 min. After fixation, the slides were rinsed briefly in 0.1% DEPC and subjected to HE staining with hematoxylin for 10 seconds and eosin for 5 seconds, with brief rinsing with 0.1% DEPC between each step. The sections were hydrated with a gradient of 70% and 100% ethanol and then air dried. All processes were performed on ice, and all reagents were prepared in 0.1% DEPC.

### 2.4. Laser Capture Microdissection

Cells in the GCL were microdissected by laser pressure catapulting (LPC) using a PALM Zeiss UV laser capture microdissection system (PALM Zeiss Microlaser Technologies, Munich, Germany) consisting of an inverted microscope with a motorized stage, an ultraviolet (UV) laser and an X-Cite 120 fluorescence illuminator (EXFO). The microdissection process was visualized with an AxioCam ICc camera coupled to a computer and was controlled by Palm RoboSoftware (Zeiss, Germany). Cells in the GCL with a large round or oval nuclei were identified as RGCs. Approximately fifty selected cells were ejected against gravity into the cap of an Eppendorf tube filled with 20 µl RNA extraction lysis buffer.

### 2.5. RNA sequencing and Gene ontology and pathway enrichment analysis

cDNA libraries were generated from the RNA samples purified from these cells using the Smart-seq2 protocol. One microliter of sample was used to determine the concentration (Qubit 3.0 fluorometer) and integrity (Agilent 2100 Bioanalyzer, Agilent High Sensitivity DNA Kit). RNA sequencing was performed using the PE100 strategy (HiSeq 2500, Illumina). Sequencing data were then analyzed. Gene expression was quantified according to RPKM (reads per kilobase per million). To analyze the biological classification of the DEGs, Gene ontology (GO) and The Kyoto Encyclopedia of Genes and Genomes (KEGG) pathway enrichment analyses were performed using DAVID. P<0.01 was used as the threshold value.

### 2.6. cDNA synthesis and real-time polymerase chain reaction (RT-PCR)

Total RNA was isolated from the GCL tissue by the “cut and LPC” method using a PALM Zeiss UV laser capture microdissection system and an RNeasy kit (Qiagen, Valencia, CA), and a portion (1 ng) of the RNA were subjected to reverse transcription (RT) with a SMARTer™ PCR cDNA Synthesis Kit (Clontech, Mountain View, CA). Then, RT-PCR analysis was performed using the TB Green® Premix Ex Taq™ Tli RNaseH Plus kit (TAKARA, Mountain View, CA). The gene-specific primer sequences were as follows: for Atf3, forward 5′-TCAGTGACAGGGCAGGAAGA −3′, reverse 5′-CCCACAGTGCAGACACCTTC −3′; for Lgals3, forward 5′-TGCCCTACGATATGCCCTTG −3′, reverse 5′-TGAAGCGGGGGTTAAAGTGG-3′; for LOC102551701, forward 5′-AACCCGATTGTGTGTTTGCG-′, reverse 5′-TTCTGTGATCGTGCTGTGCT-3′; for Plaur, forward 5′-GCGGCCGCGAAGAACC-3′, reverse 5′-CTTCCCATTCCCGAAGCACC-3′; for Tmem140, forward 5′-ATGGGATGGCAAGGGTTCAG-3′, reverse 5′-TGGGTTCCTTGACACCACAG-3′; for Maml1, forward 5′-CGGACATCTCCATGATCCAGT-3′, reverse 5′-GAAGCAGGAGGAAGCCCATT-3′.

### 2.7. Immunohistochemistry

Rats were sacrificed and perfused intracardially with 4% paraformaldehyde (PFA) in phosphate buffer saline (PBS). The eyes were removed and immersed in 4% PFA in PBS for 24 h at 4°C and then in 10, 20, and 30% sucrose solution in PBS for 24 h at 4°C and embedded using optimal cutting temperature embedding medium (SAKURA Tissue-Tek® O.C.T. Compound) by rapid freezing under crushed dry ice, after which they were stored at −80°C. After embedding, the eyes were sectioned with a CM1950 cryostat microtome (Leica Microsystems Inc, Bannockburn, IL) at −22°C at a thickness of 15 mm. The parasagittal eye sections were stored at 22°C before immunocytochemistry, and eye sections with a visible optic nerve head were chosen.

Tissues were washed in PBS (3×5 min) and then incubated in blocking solution (5% donkey serum, 0.5% Triton X-100, and 1% bovine serum albumin in PBS) for 1 h at room temperature (RT), followed by incubation with primary antibodies overnight at 4°C. The primary antibodies used were rabbit anti-Atf3 (1:500; Abcam) and rabbit anti–Maml1 (1:500; Abcam). The next day, tissues were washed in PBS (3×5 min) and incubated with secondary antibodies conjugated to Alexa Fluor 488 (1:400; goat anti–rabbit; Life Technologies) or Alexa Fluor 594 (1:400; goat anti–rabbit; Life Technologies) for 1 h at RT. Retinas were incubated with the nuclear dye DAPI for 20 min and washed in PBS (3×5 min) before microscopic analysis.

All sections were observed under a fluorescence microscope (Leica DFC 7000T), and photomicrograph images were captured at 200× magnification.

### 2.8. Statistical analysis

Continuous variables were compared using the t test after the normal distribution test. Categorical variables were compared using χ2 analysis. Ranked variables were compared using non-parametric tests along with the Mann–Whitney U test. Differences were considered statistically significant when p ≤ 0.05.

## 3. Results

### 3.1. Workflow used for laser capture microdissection and single-cell RNA sequencing

Our protocol was divided into two parts: optimization of the LCM protocol and determination of the ONC-related early mRNA alterations in RGCs.

In part one, corneas and lenses were removed from normal rat eyes and prepared for sectioning. Glass slides and PEN membrane-coated slides were used, and tissue sections with a thickness of 20 µm or 30 μm were used. The effects of section thickness and slide type on the tissue capture success and RNA quality of RGCs after LPC using the PALM Zeiss UV LCM system were analyzed (Fig. 1A and C).

**Fig.1.**
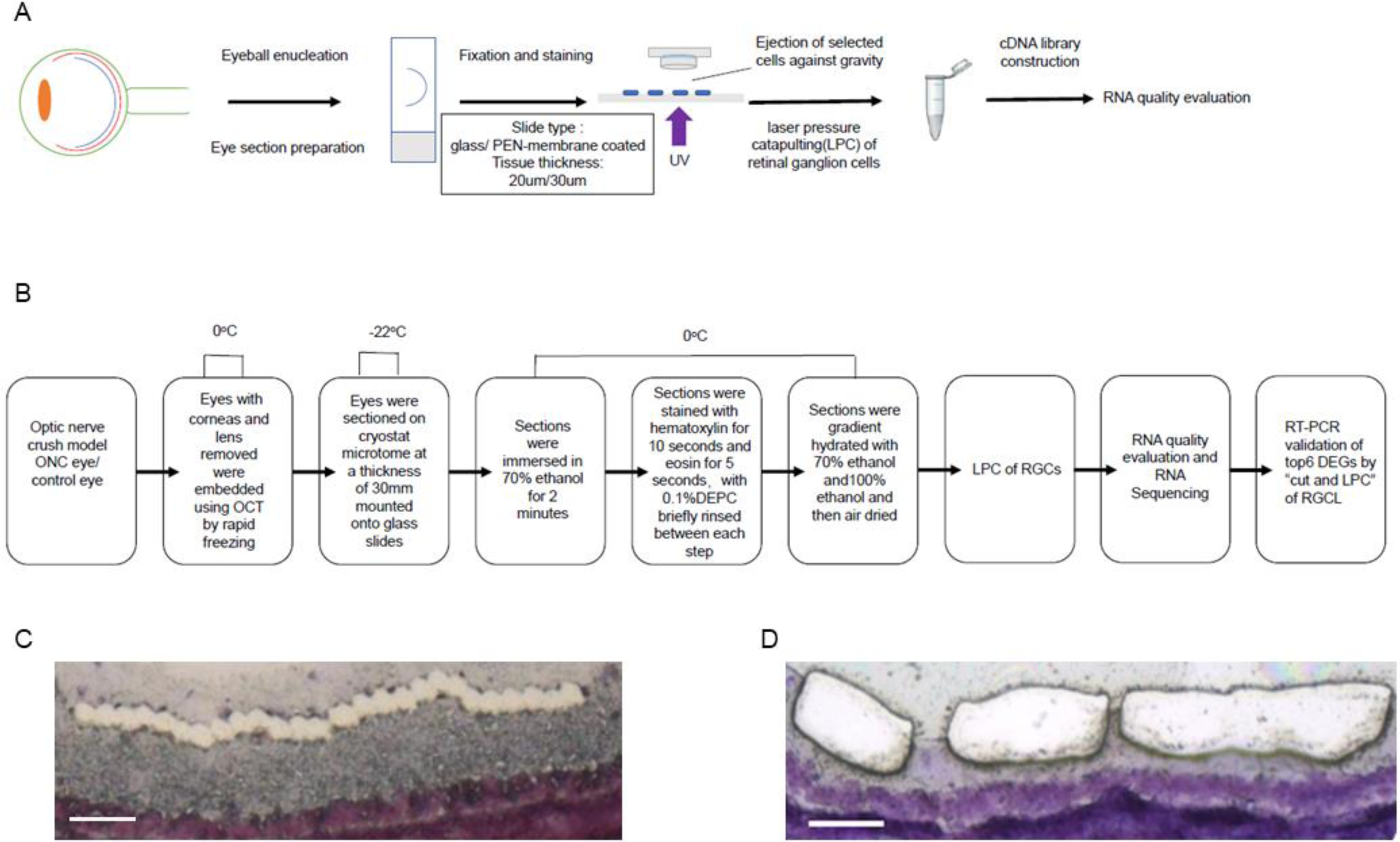
Workflow of laser capture microdissection-based sequencing. (A) Workflow of optimizing the LCM protocol,(B) Workflow of the study of early mRNA alterations in RGCs after ONC,(C) LPC of RGCs for mRNA sequencing (bar=50 µm), (D) “Cut and LPC” method for RGCL for RT-PCR (bar=50 µm)

In part two, the optimized LCM protocol (30 μm-thick tissue sections mounted on glass slides) was used to explore ONC-related early mRNA alterations in RGCs. Candidate genes were validated by RT-PCR of GCL tissue dissected by the “cut and LPC” method using the same LCM system describe above. (Fig. 1B and D)

### 3.2. Effect of section thickness and slide type on tissue capture success, RNA yield, and integrity after LCM

Retinal sections with two different thicknesses (20 or 30 µm) that were mounted onto two different types of slides (glass or PEN-membrane coated) were used to determine the best tissue preparation conditions for LCM. The capture success was calculated by dividing the dissected cell number (after the LCM process) by the selected cell number (before the LCM process). Both the 20 µm-thick and 30 µm-thick sections mounted on PEN membrane-coated slides led to poor capture success (16% vs 20%, p>0.05) since no more than 20% of the selected cells were dissected. And sections mounted on glass slides showed good capture success (both 100%). Regarding the RNA integrity, all the samples from 30 µm-thick sections on glass slides met the demands for library construction, while only 75% of the samples from 20 µm-thick sections were acceptable (p>0.05). Regarding the RNA yield, the 20 µm-thick sections showed a yield of 1.52±0.25 ng/µl, and the 30 µm-thick sections showed a yield of 2.29±0.31 ng/µl (p<0.05, Fig. 2A and B). These results showed that PEN membrane-coated slides did not lead to good cell capture. Thirty-micrometer-thick glass sections had higher RNA yield than 20-µm-thick glass sections, with both samples resulting in good RNA integrity for further RNA sequencing analysis.

**Fig.2.**
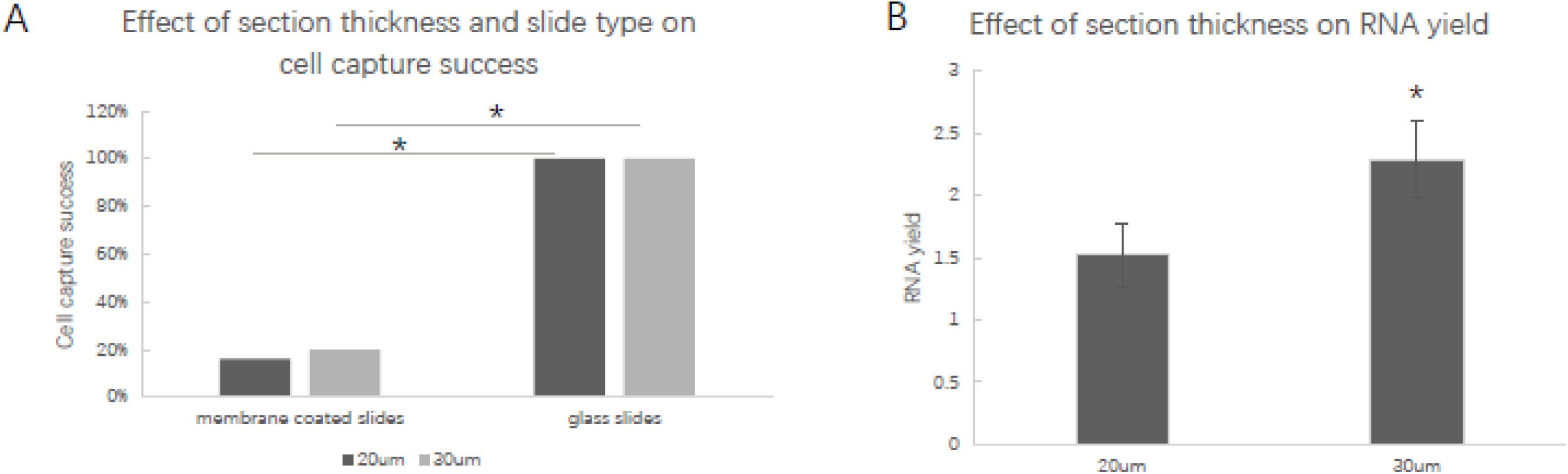
Effect of section thickness and slide type on cell capture success, RNA yield, and integrity after LCM. (A) Effect of section thickness and slide type on cell capture success. Both 20 µm-thick and 30 µm-thick sections mounted on PEN membrane-coated slides led to poor capture success (16% vs 20%, p>0.05). Sections mounted on glass slides led to good capture success (both 100%). (B) Effect of section thickness on RNA yield. Twenty-micrometer-thick sections showed a yield of 1.52±0.25 ng/µl, and 30-µm-thick sections showed a yield of 2.29±0.31 ng/µl (p<0.05).

### 3.3. RNA sequencing analysis and identification of differentially expressed genes (DEGs) 3 days postcrush

The RNA from dissected cells was then amplified and used to generate cDNA libraries using the Smart-seq2 protocol. After RNA sequencing employing the PE100 strategy (HiSeq 2500, Illumina), the dissected cells were identified by cell markers. RGC markers such as Pou4 (Brn3), Rbpms and Slc17ab were shown to be expressed with high FPKM values (Fig. 3A). Astroglial markers such as S100, GFAP, MAP2, and ALDH1L1 were shown to be expressed with low FPKM values. The horizontal cell marker 8A-1, microglia marker Iba-1, and amacrine cell marker HPC-1 were not found in our results.

**Fig.3.**
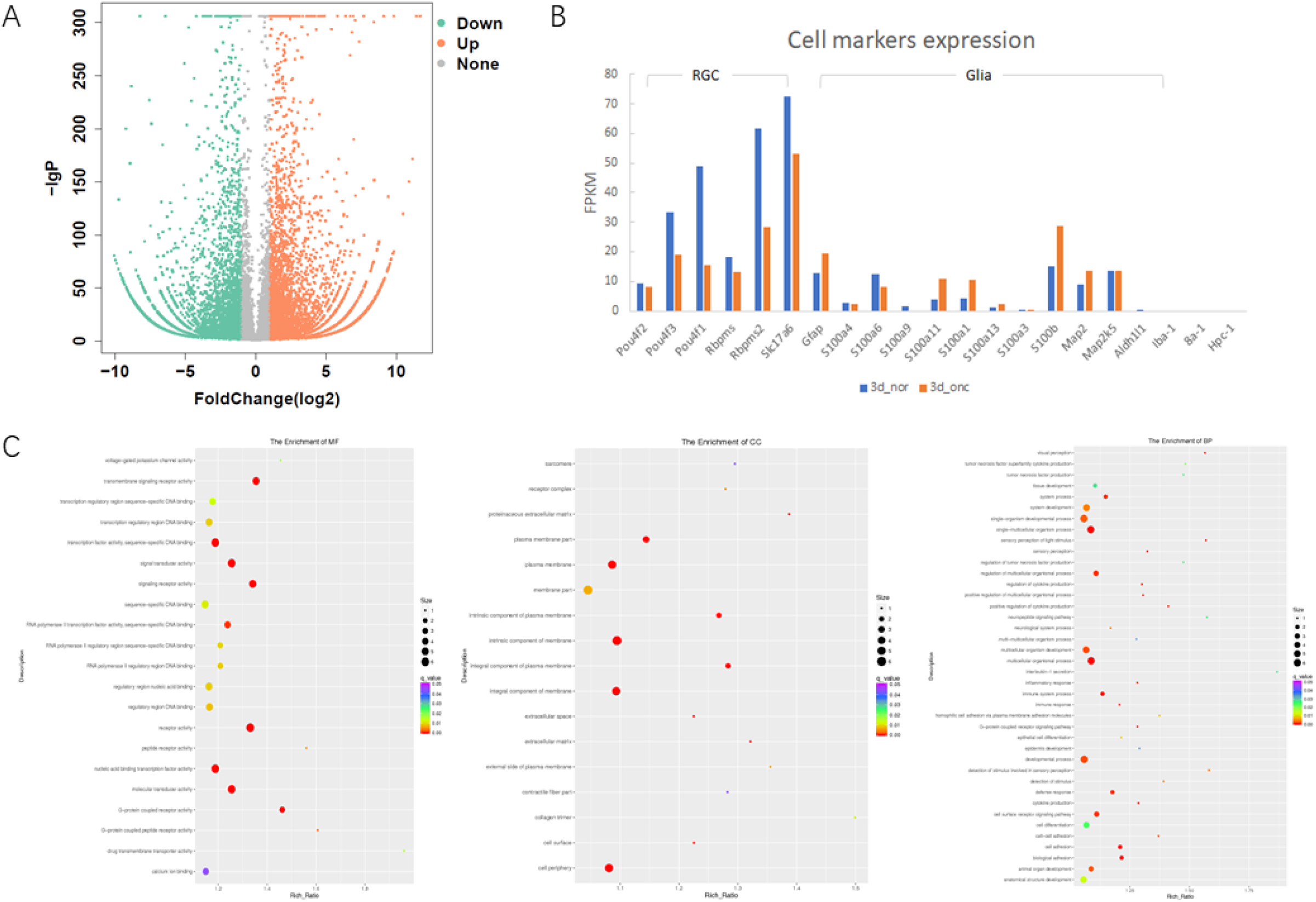
Analysis of scRNA-seq results. (A) Biomarkers of GCL cells. RGC markers were identified to be expressed with high FPKM values, while other markers were identified with low FPKM values. (B) Volcano plot of DEGs. Genes with a fold change ≥ 2 and a q value < 0.05 were chosen. A total of 8744 DEGs were included, with 4411 upregulated genes and 4333 downregulated genes. (C) GO analyses of DEGs. DEGs were enriched in BP, MF and CC.

Comparison of the differential expression profiles of mRNAs from ONC and control RGCs was performed after normalization of the raw data and application of DESeq2. Genes with a fold change ≥ 2 and a q value < 0.05 were chosen. A total of 8744 differentially expressed genes (DEGs) were included, with 4411 upregulated genes and 4333 downregulated genes (Fig. 3B). The candidate genes were chosen from among DEGs with a |log2Ratio| ≥ 10, including five genes that were unexpressed and one gene that was downregulated in ONC RGCs compared with that in control RGCs. (Table. 1)

**Table.1.** Differentially expressed genes with a |log2Ratio|⩾10 in RGCs after ONC.

| Gene | Log2 fold change | padj | Up/down | Gene name | Full name |
| --- | --- | --- | --- | --- | --- |
| gene27639 | 11.70331 | 0 | up | Atf3 | activating transcription factor 3 |
| gene29150 | 11.4495 | 0 | up | Lgals3 | lectin, galactoside-binding, soluble, 3 |
| gene35168 | 11.15631 | 2.35E-172 | up | LOC102551701 |  |
| gene1274 | 10.90482 | 3.32E-151 | up | Plaur | Plasminogen Activator, Urokinase Receptor |
| gene10710 | 10.47381 | 1.28E-120 | up | Tmem140 | transmembrane protein 140 |
| gene22509 | -10.0667 | 2.07E-81 | down | Maml1 | Mastermind-like 1 |

### 3.4. GO and KEGG enrichment analyses of the DEGs

To analyze the biological classification of the DEGs, functional and pathway enrichment analyses were performed using DAVID. The GO analysis results showed that the DEGs related to biological processes (BP) were significantly enriched in multicellular organismal process, single-multicellular organism process, and biological adhesion, etc. DEGs with changes in molecular function (MF) were mainly enriched in receptor activity, signaling receptor activity, and transmembrane signaling receptor activity, etc. DEGs with changes in cell components (CC) were mainly enriched in integral components of the plasma membrane, intrinsic components of the plasma membrane, and intrinsic components of the membrane, etc. (Fig. 3C)

An additional functional analysis of mRNAs based on KEGG pathways showed that the neuroactive ligand−receptor interaction pathway was significant changed in ONC versus that in normal RGC. There were 103 genes in the pathway that were changed, including 38 upregulated and 65 downregulated genes.

### 3.5. RT-PCR gene expression analysis

GCL was microdissected by the “cut and LPC” method using the PALM MicroBeam system, and the RNA was extracted using a column-based method. Total RNA was subjected to reverse transcription (RT) with a SMARTer™ PCR cDNA Synthesis Kit and then used for RT-PCR analysis. Atf3(activating transcription factor 3), Lgals3 (lectin, galactoside-binding, soluble, 3), LOC102551701, Plaur (plasminogen activator, urokinase receptor) and Tmem140 (transmembrane protein 140) levels were unregulated in the cells from ONC eyes compared to the mRNA levels in the control eyes (p<0.01), while the Maml1 (Mastermind-like 1) mRNA level was decreased in the ONC eyes compared to that in the control eyes (p<0.01) (Fig. 4A).

**Fig.4.**
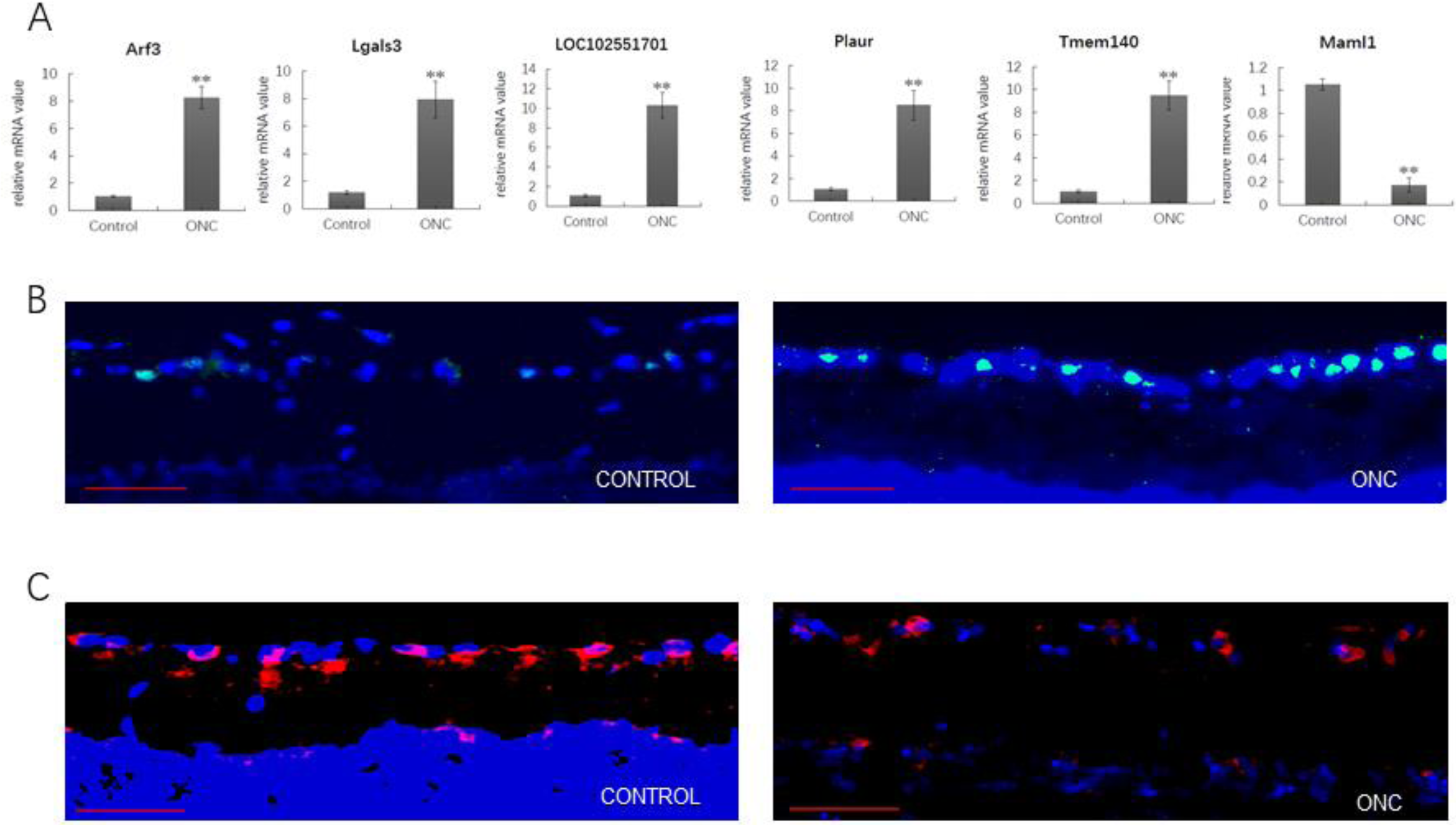
Verification of candidate genes. (A) RT-PCR of DEGs with a |log2Ratio|≥10. Atf3, Lgals3, LOC102551701, Plaur and Tmem140 levels were found to be upregulated, while Maml1 mRNA was found to be decreased in the ONC eyes compared to the control eyes (p<0.01) (B) Immunohistochemistry of ATF3. In the control retina, there was low Atf3 immunoreactivity, while in the ONC retina, some cells in the GCL expressed Atf3 at a high level (green-ATF3, blue-DAPI, bar=200 µm). (C) Immunohistochemistry of Maml1. In control retinas, there was high Maml1 immunoreactivity, while in ONC retinas, Maml1 immunoreactivity was low in the GCL (red-Maml1, blue-DAPI, bar=200 µm).

### 3.6. Immunohistochemistry of Atf3 and Maml1

Retinal immunostaining of the Atf3 and Maml1 proteins confirmed the apparent changes in protein expression after ONC. In the control retinas, there was low Atf3 immunoreactivity and obvious Maml1 immunoreactivity throughout all retinal layers. In the ONC retinas, some cells in the GCL expressed Atf3 at a high level in the GCL, and Maml1 expression was hardly observed in the GCL (Fig. 4B and C).

## 4. Discussion

LCM is often used to study the expression of both minimally or abundantly expressed genes in different brain regions[5-9] and occasionally in retinas[10-12]. In our study, we adopted an optimal LCM protocol with single-cell sequencing to examine the early mRNA changes in RGCs in an optic nerve crush model.

During the LCM process, capture success depends on the tissue section thickness and the slide type. A previous study showed that 10 µm-thick tissue sections were not adequate for the capture of microdissected tissue and that the resulting RNA was totally degraded[5]. In our study, we found that 30 µm-thick tissue sections had high capture success and RNA yield when using the PALM system. However, this might not be applicable to another LCM system, such as the Arcturus LCM system as it possesses a low-energy infrared laser which can not catapult thicker tissue sections [13].

It was reported that membrane-coated slides made it possible to capture larger tissue areas with lower laser intensity and time compared with glass slides[14]. However, others reported that tissue sections mounted on glass slides led to higher capture success and RNA yield compared with sections mounted on PEN membrane-coated slides[5]. The increase in adhesion between tissue and PEN-membrane slides made it difficult to catapult the tissue, which was also observed in our study.

We compared our sequencing results with the currently published 10x scRNA-seq results for RGCs[2] and found that there were a large number of ganglion cell markers observed in our sequencing data, such as Pou4, Rbpms and Slc17ab, which meant our sequencing process could obtain ganglion cells accurately and was more economical and thus more suitable for low-throughput PCR to detect specific genes in small samples.

We also found a large number of DEGs 3 days after ONC, including Atf3, Lgals3, LOC102551701, Plaur, Tmem140 and Maml1. Among those. Atf3 was previously found to be significantly changed in ONC GCL[12]. This result was in accord with our results. Atf3 was reported to be involved in axonal regeneration, as Atf3 expression was activated following crush, and axonal regeneration was enhanced in various types of neurons as a result[15] [16] [17]. This implied that axonal regeneration started in the early stage after ONC.

Lgals3 is a multifunctional protein that participates in mediating inflammatory reactions and cell proliferation, adhesion, migration, and apoptosis. The expression of Lgals3 was reported in astrocytes and microglia induced by neuroinflammation [18] [19] and was involved in autophagy in the central nervous system (CNS)[20]. The increased in Lgals3 after ONC in our results might be due to its expression by activated astrocytes in the RGC layer.

Plaur, the receptor for the plasminogen activator of the urokinase type (uPAR), is involved in many physiological and pathological processes and participates in the development, function and pathology of the central nervous system[21] The uPAR leads to activation of the Rho family small GTPase Rac1 and induced axonal regeneration in the CNS.[22] We also suggest the increased Plaur levels according to our results could lead to Rac1-induced axonal regeneration after ONC, which needs further study.

Another interesting candidate was Maml1. The close correlation of the spatial and temporal expression of Maml1 in the CNS during early development indicates the role of the Maml1 gene in neurogenesis. [23] MAML1 is also a novel modulator of NF-kappa B signaling and regulates cellular survival.[24] Hence, the decrease in Maml1 in the early stage after ONC might reflect the insufficient intrinsic growth state of mature neurons.

There are some limitations of our study. First, the results were based on microtranscriptome sequencing of a small number of cells, which might not be adequate to reflect the whole RGC population. Second, since RGCs were identified based on HE staining, there might be small numbers of glial cells or retinal intermediate neuron cells mixed in the sample. Further studies and improved protocols are needed to discover the mechanism involved in changes after ONC.

## Abbreviations

RGC: retinal ganglion cell
GCL: ganglion cell layer
ONC: optic nerve crush
LCM: laser capture microdissection
scRNA-seq: single-cell RNA sequencing
LPC: laser pressure catapulting
PFA: paraformaldehyde
PBS: phosphate buffer saline
uPAR: plasminogen activator of the urokinase type
LRP1: lipoprotein receptor-related protein-1
Atf3: activating transcription factor 3
Lgals3: lectin, galactoside-binding, soluble, 3
Plaur: Plasminogen Activator, Urokinase Receptor
Tmem140: transmembrane protein 140
Maml1: Mastermind-like 1

## Acknowledgements

This research was funded by the China National Key Research and Development Program Stem Cell and Translational Research Key Projects (2018YFA0108301), National Natural Science Foundation of China (31971109, 31471390), Shanghai Rising-Star Program (17QA1405400) and Shanghai Health and family planning system program (2017YQ028).

We also thank for Prof. Jiajun Xu’s critical proofreading of the manuscript. Y.W., X.D.S., W.S. and D.P. designed the research. D.Y.P., M.Q.X., X.C.,Y.B.F.,Y.H.F and M.X. performed the experiments. D.Y.P., M.Q.X., X.C., and Y.W. analyzed the data, and D.Y.P. and Y.W. wrote the manuscript.

